# “What” and “when” predictions jointly modulate speech processing

**DOI:** 10.1101/2024.05.10.593519

**Authors:** Ryszard Auksztulewicz, Ozan Bahattin Ödül, Saskia Helbling, Ana Böke, Drew Cappotto, Dan Luo, Jan Schnupp, Lucía Melloni

**Affiliations:** Centre for Cognitive Neuroscience Berlin, Freie Universität Berlin, Germany; Department of Brain and behavioral Sciences, Università di Pavia, Italy; Ernst Strungmann Institute, Frankfurt am Main, Germany; Ear Institute, University College London, UK; Department of Otorhinolaryngology, Perelman School of Medicine, University of Pennsylvania, USA; Department of Neuroscience, City University of Hong Kong, Hong Kong SAR; Research Group Neural Circuits, Consciousness and Cognition; Max Planck Institute for Empirical Aesthetics, Frankfurt am Main, Germany

## Abstract

Adaptive behavior rests on forming predictions based on previous statistical regularities encountered in the environment. Such regularities pertain not only to the contents of the stimuli (“what”) but also their timing (“when”), and both interactively modulate sensory processing. In speech streams, predictions can be formed at multiple hierarchical levels, both in terms of contents (e.g. single syllables vs. words) and timing (e.g., faster vs. slower time scales). Whether and how these hierarchies map onto each other in terms of integrating “what” and “when” predictions remains unknown. Under one hypothesis neural hierarchies may link “what” and “when” predictions within sensory processing areas: with lower cortical regions mediating interactions for smaller units e.g., syllables, and higher cortical areas mediating interactions for larger units e.g., words. Alternatively, interactions between “what” and “when” predictions might rest on a generic, sensory-independent mechanism, mediated by common attention-related (e.g., frontoparietal) networks. To address those questions, we manipulated “what” and “when” predictions at two levels – single syllables and disyllabic pseudowords – while recording neural activity using magnetoencephalography (MEG) in healthy volunteers (N=22). We studied how syllable and/or word deviants are modulated by “when” predictability, both analyzing event-related fields and using source reconstruction and dynamic causal modeling to explain the observed effects in terms of the underlying effective connectivity. “When” predictions modulated “what” mismatch responses in a specific way with regards to speech hierarchy, such that mismatch responses to deviant words (vs. syllables) were amplified by temporal predictions at a slower (vs. faster) time scale. However, these modulations were source-localized to a shared network of cortical regions, including frontal and parietal sources. Effective connectivity analysis showed that, while mismatch responses to violations of “what” predictions modulated connectivity between regions, the integration of “what” and “when” predictions selectively modulated connectivity within regions, consistent with gain effects. These results suggest that the brain integrates “what” and “when” predictions that are congruent with respect to their hierarchical level, but this integration is mediated by a shared and distributed cortical network. This contrasts with recent studies indicating separable networks for different levels of hierarchical speech processing.

## 1. Introduction

Speech comprehension is grounded in auditory predictions, enabling the brain to anticipate future acoustic events and facilitate efficient processing (Caucheteux et al., 2023; Hickok, 2012; Hovsepyan et al., 2020; Mai and Wang, 2023; Poeppel and Assaneo, 2020). When these predictions are accurate, they can significantly reduce the amount of bottom-up processing required, allowing the brain to optimize for other aspects of language comprehension (Federmeier, 2007; Hakonen et al., 2017; Ryskin and Nieuwland, 2023; Sohoglu and Davis, 2016). Conversely, when predictions are inaccurate or conflicting, they draw cognitive resources to resolve these inconsistencies, leading to error signals or processing delays (Blank et al., 2018; Cope et al., 2017). Studies investigating the neural correlates of predictions in speech processing have shown that several brain regions, including the superior temporal gyrus (STG), superior parietal lobule (SPL), and prefrontal cortex, are implicated in the processing of predictive signals (Caucheteux et al., 2023; Shain et al., 2020). The timing of these neural responses suggests that predictions are generated at multiple levels of the auditory hierarchy, from low-level acoustic properties to high-level semantic representations (Caucheteux et al., 2023; Schmitt et al., 2021).

Predictions in speech processing encompass multiple stimulus features, such as the contents (“what”) and timing (“when”) of speech units (Gómez Varela et al., 2024). When unexpected auditory contents are encountered even in artificial speech, those prediction errors are reflected in neural activity (Ylinen et al., 2016). Studies have shown that these prediction errors typically increase activity in the inferior frontal gyrus (IFG) (Petersson et al., 2012; Wilson et al., 2015), as well as in the superior temporal cortex (Ling et al., 2022), among other regions. Importantly, “what” predictions can span multiple levels, ranging from phonemes to syllables, words, and longer phrases (Heilbron et al., 2022; Su et al., 2023). This suggests that “what” predictions may be hierarchically organized, with higher-level predictions guiding the interpretation of lower-level information (Heilbron et al., 2022).

Similarly, “when” predictions can operate across different time scales, from phoneme onsets to phrase boundaries and beyond (Ding et al., 2016; Donhauser and Baillet, 2020; Schmitt et al., 2021). Many speech sequences exhibit a consistent temporal pattern that can entrain neural oscillations (Ding et al., 2016). This entrainment, also known as phase locking, is reflected in phasically increased neural activity in regions involved in auditory processing (Obleser and Kayser, 2019) including the STG (Keitel et al., 2017), although it is debated to what degree increased phase-locking reflects a true oscillatory mechanism rather than a series of evoked responses (Doelling et al., 2019; Oganian et al., 2023). In contrast, jittered speech sequences, which lack a consistent temporal pattern, exhibit weaker neural entrainment (Klimovich-Gray et al., 2021). Research suggests that the brain’s ability to entrain to rhythmic patterns plays a role in processing and understanding speech, particularly in situations where the temporal structure of the speech signal is predictable (Kösem et al., 2018; Riecke et al., 2018; Zoefel et al., 2018).

While “what” predictions and “when” predictions are both essential for efficient speech comprehension, they are likely subserved by different mechanisms (Arnal and Giraud, 2012; Auksztulewicz et al., 2018). In a study combining invasive electrophysiological measurements with computational modeling, we proposed that the most likely mechanisms of “what” predictions of auditory events rest on modulated connectivity between sensory, prefrontal, and premotor regions. In contrast, we proposed that “when” predictions are predominantly linked to gain modulation in sensory regions independently of activity in other cortical regions (Auksztulewicz et al., 2018). Increased gain due to (especially rhythmic) “when” predictions (Auksztulewicz et al., 2019) may also result in strong interactions between “what” and “when” predictions, such that mismatch responses evoked by mispredicted auditory stimuli are amplified when these stimuli are presented in rhythmic (temporally predictable) sequences (Jalewa et al., 2021; Lumaca et al., 2019; Takegata and Morotomi, 1999; Todd et al., 2018; Yabe et al., 1997).

Given the hierarchical nature of speech signals, the question remains: how do “what” and “when” predictions interact at the appropriate levels of the speech hierarchy? One plausible hypothesis maps the hierarchies of “what” and “when” predictions onto neural hierarchies, such that the interactive effects of “what” predictions for single chunks (e.g., syllable) and “when” predictions for faster time scales (e.g., syllable onsets) are subserved by hierarchically lower cortical regions involved in syllable processing, such as the STG (Oganian and Chang, 2019). Conversely, interactions between “what” predictions for longer segments (e.g., words) and slower “when” predictions (e.g., word onsets) may instead be subserved by hierarchically higher cortical regions involved in supra-syllabic word processing, such as frontal regions (Rimmele et al., 2023). Yet, interactions between “what” and “when” predictions may not need to occur within the sensory processing hierarchy, and instead might rest on sensory-independent, generic mechanisms regardless of their hierarchical level in terms of speech contents and timing. Such generic mechanisms may rely on attention-related brain regions e.g., the left parietal cortex, previously shown to be involved in integrating content-based and time-based information in speech (Orpella et al., 2020).

To disambiguate these hypotheses, the current study investigates the neural correlates of “what” and “when” predictions across different hierarchical levels of artificial speech stimuli. By separately manipulating the predictability of linguistic content and temporal alignment at multiple levels, we aim to determine whether and how these predictions interact in terms of neural responses evoked by speech stimuli (as measured using magnetoencephalography; MEG). To that end, we first quantify the extent of phase-locking of MEG responses to syllables and words across different types of “when” predictions, and then we use source reconstruction and computational modeling of the evoked responses to infer the putative mechanisms underlying interactive predictive speech processing across the cortical hierarchy.

## 2. Results

The aim of the analysis was twofold. First, to examine the impact of “when” predictions on responses to syllables and on neural entrainment at lower vs. higher timescales. Second, to assess how these “when” predictions modulate neural signatures associated with lower and higher-level “what” predictions, specifically the mismatch responses (MMRs). The latter analysis focused on testing interactions between “what” and “when” predictions, with a specific emphasis on testing whether MMRs exhibit contextually specific modulation based on temporal predictability. For instance, the analysis probed whether faster (vs. slower) “when” predictions selectively influence MMRs in response to violations of “what” predictions concerning syllables (vs. words). To draw inferences on the putative mechanisms underlying interactions between “what” and “when” predictions observed at the sensor level, we performed source reconstruction and model-driven data analysis techniques (dynamic causal modeling)

### 2.1. Behavioral results

Prior to MEG measurements, participants (N=22) implicitly learned 6 disyllabic pseudowords (“tupi”, “robi”, “daku”, “gola”, “zufa”, “somi”; Fig. 1A) in a two-minute block of passive exposure (see Methods for more details). We first tested if participants implicitly learned the pseudowords. In line with that hypothesis, we observed that pseudoword discrimination following initial training far exceeded chance level, reaching 71.35% (SEM: 1.81%; one-sample t-test against chance level: t_20_ = 11.77, p < 0.001; Fig. 2A).

**Figure 1.**
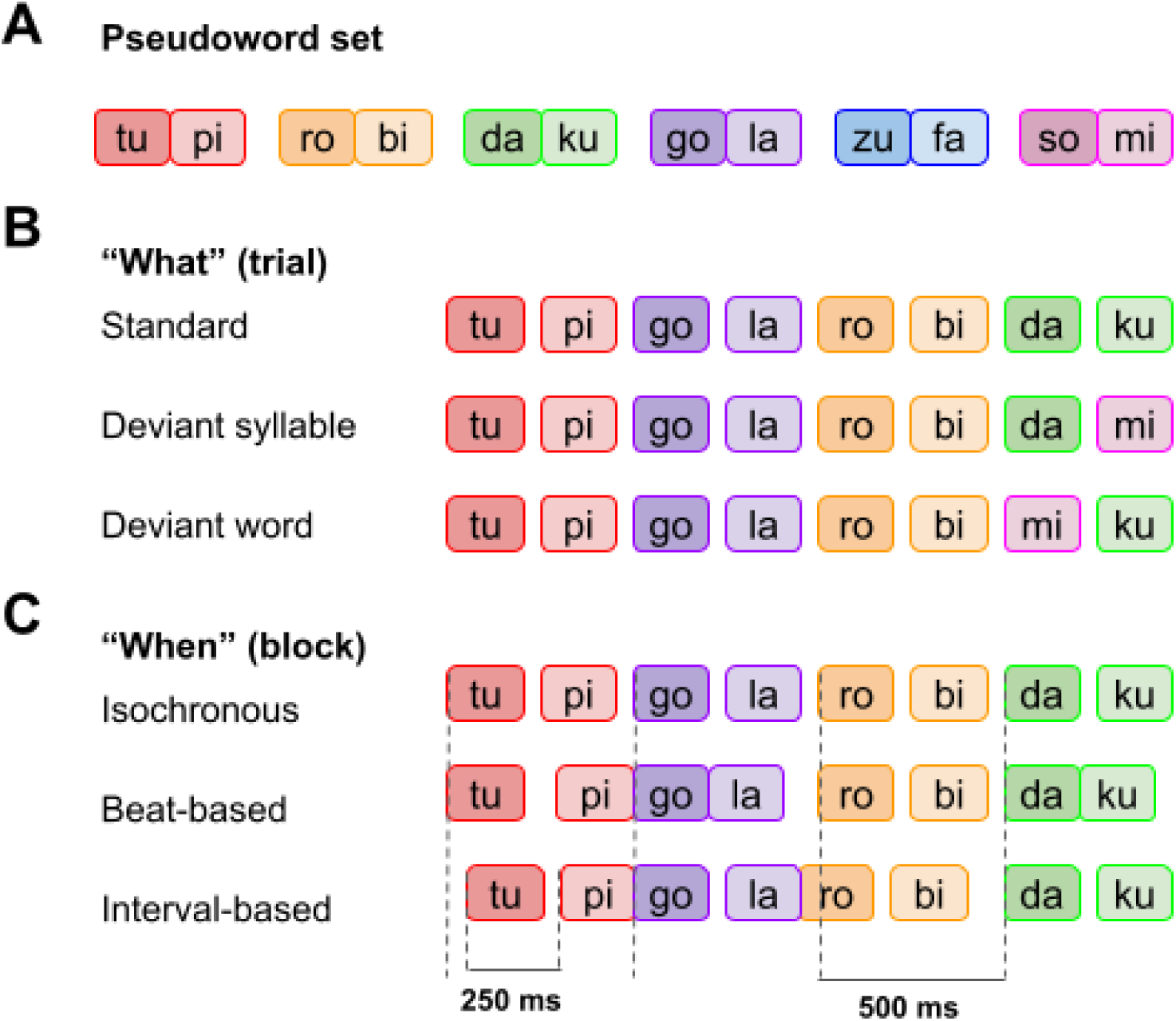
Experimental paradigm. **(A)** Participants listened to sequences of six pseudowords to which they had been passively exposed in a training session. **(B)** After participants had implicitly learned the six pseudowords, they engaged in a syllable repetition detection task. During the task, “what” predictions were manipulating across three experimental conditions (at a single trial level): pseudoword sequences could be composed of only legal pseudowords (“standard” trials), or they could contain a pseudoword with a deviant word-final syllable (“deviant syllable” trials, whereby the pseudoword starts in an expected manner but ends with a violation), or they could contain a pseudoword with a deviant word-initial syllable substituted from a final syllable of another pseudoword (“deviant word” trials, whereby the pseudoword starts with a violation). **(C)** Sequences were blocked into three temporal conditions: an “isochronous” condition, in which ISI between all syllables was fixed at 0.25 s; a “beat-based” condition, in which the ISI between pseudoword onsets was fixed at 0.5 s but the ISI between the initial and final syllables of each pseudoword was jittered, such that only the timing of the deviant word (but not of the deviant syllable) could be predicted; and an “interval-based” condition, in which the ISI between pseudoword onsets was jittered but the ISI between the initial and final syllables of each pseudoword was fixed at 0.25 s, such that only the timing of the deviant syllable (but not of the word) could be predicted.

**Figure 2.**
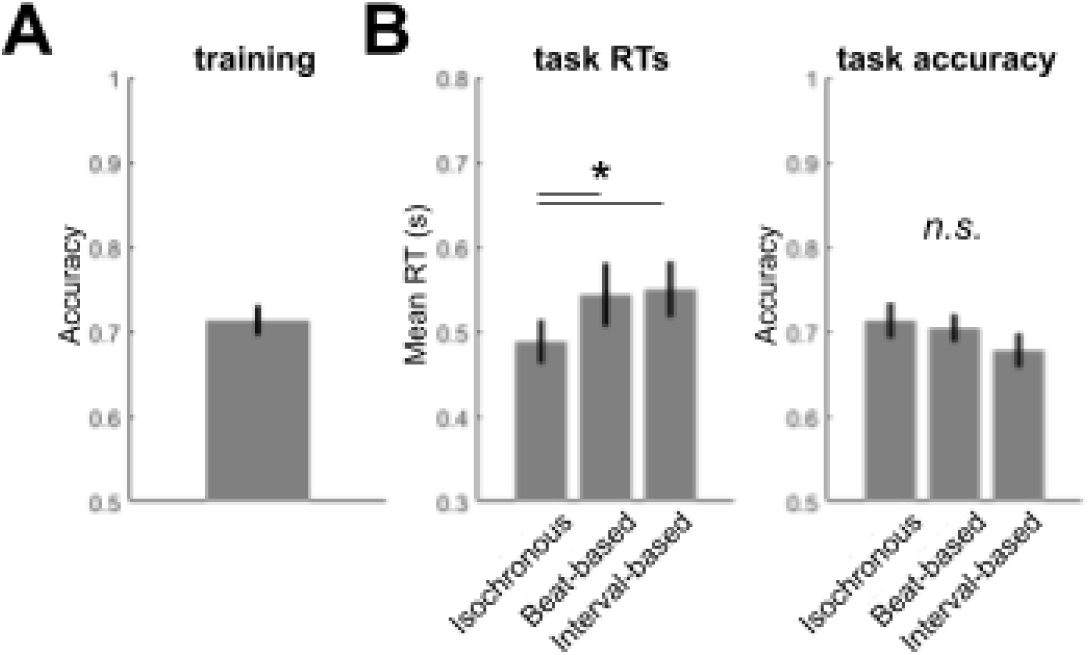
**Behavioral results. (A)**: Accuracy in the training session; **(B)** Left panel: reaction times during the repetition detection task, right panel: accuracy during the repetition detection task. Error bars denote SEM across participants. Asterisks denote p < 0.05.

After participants had learned the set of pseudowords, they listened to a continuous stream of syllables, engaging in a syllable repetition detection task while MEG was measured. The stimulus sequences were independently manipulated with respect to “what” and “when” predictions at two different levels: single syllables and disyllabic pseudowords. “What” prediction manipulations had three levels (Fig. 1B): (1) standard sequences (66.6%), in which only “correct” pseudowords were used; (2) “deviant syllable” sequences (13.3%), in which the final syllable of the sequence was replaced with a syllable belonging to a different pseudoword; (3) “deviant word” sequences (13.3%), in which the penultimate syllable of the sequence was replaced with a syllable which should be a final syllable of a different pseudoword, resulting in an unpredictable disyllabic pseudoword. The remaining 7% trials contained task-related repetitions that were discarded from MEG analysis. Independently, “when” prediction manipulations also had three levels (Fig. 1C): (1) isochronous sequences (33.3%), in which the SOA between consecutive syllables was fixed at 250 ms; (2) “beat-based” sequences (33.3%), in which the timing of word onsets was predictable but the timing of word-final syllables was unpredictable; and (3) “interval-based” sequences (33.3%), in which the timing of word onsets was unpredictable but the timing of word-final syllables was predictable.

Participants performed well and comparable across conditions in the syllable detection task (F_2,40_ = 1.730, p = 0.190; Fig. 2B). Yet, we observed a clear effect of temporal predictability in the RTs (F_2,40_ = 5.188, p = 0.009): in the isochronous condition participants were faster (mean ± SEM: 488 ± 26 ms, after exponentiating log RTs) than both in the interval-based (mean ± SEM: 550 ± 34 ms; t_20_ = -2.475, p = 0.022, FDR-corrected) and beat-based conditions (mean ± SEM: 544 ± 38 ms; t_20_ = -3.429, p = 0.003, FDR-corrected) (Fig. 2B). This result indicates that participants capitalized on the temporal predictability, adjusting their responses based on the available information.

### 2.2. MEG results: phase locking

The stimulus spectrum, quantified as inter-trial phase coherence (ITPC) of the sound envelope across 80 unique sequences per condition, showed pronounced differences between the three “when” conditions (Fig. 3AC). Specifically, (1) in isochronous sequences, both a prominent syllable-rate (4 Hz) and a word-rate (2 Hz) peak were found; (2) in beat-based sequences, the word-rate (2 Hz) peak was largely preserved, and a syllable-rate (4 Hz) peak was relatively weaker but still present; (3) in interval-based sequences, both peaks were relatively weaker compared to the other conditions. All pairwise differences for the 2-Hz rate as well as for the 4-Hz rate were significant (all p < 0.001, all t_21_ > 4.95).

**Figure 3.**
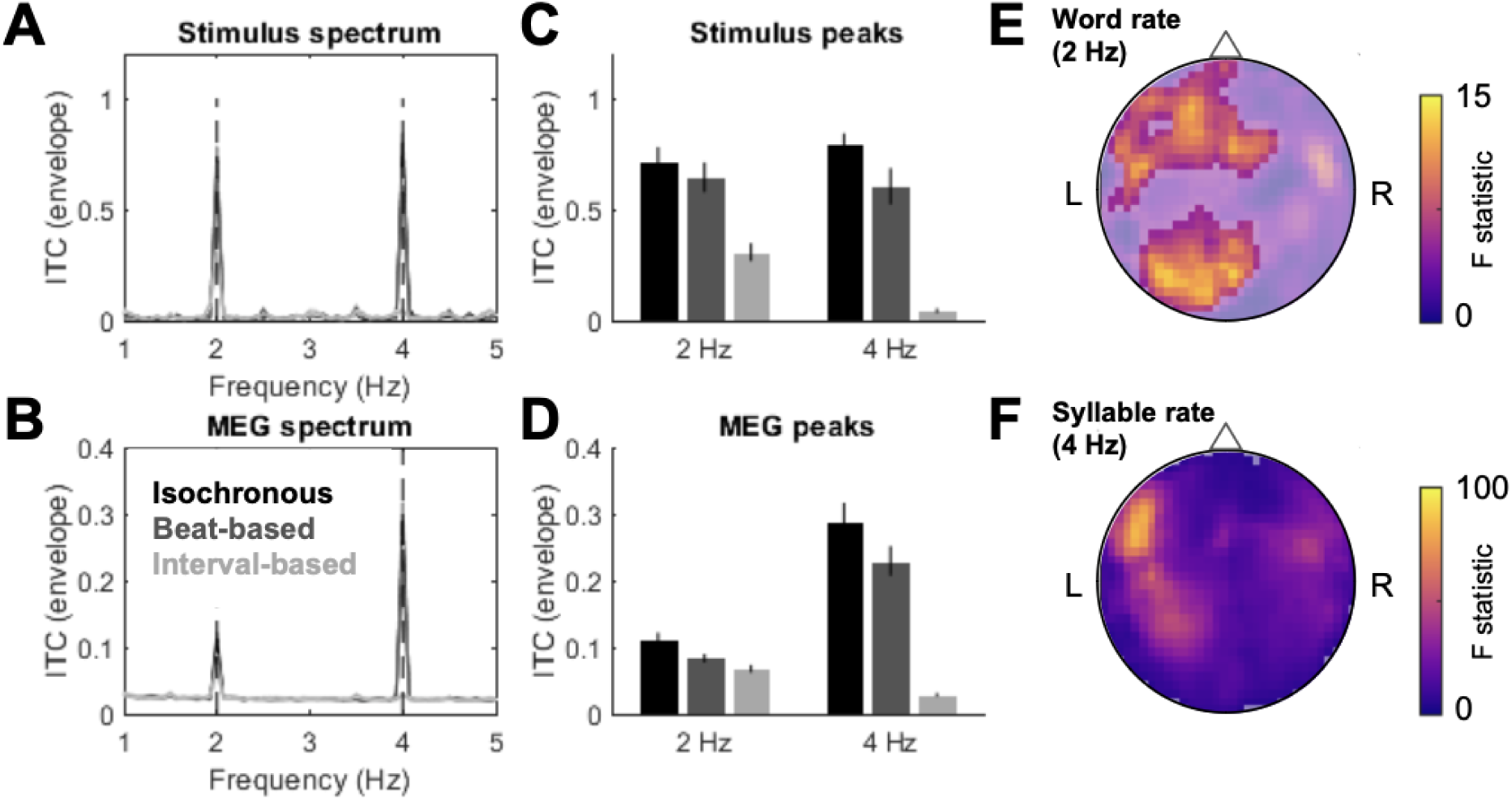
Phase locking results. **(A)** Stimulus spectrum. Prominent peaks are observed at the 2 Hz (pseudoword rate) and 4 Hz (syllable rate). **(B)** MEG spectrum, averaged across channels. Similar peaks are observed as in the stimulus spectrum. **(C)** Bar plots of 2-Hz and 4-Hz peaks based on the stimulus spectrum, showing differences between conditions. All pairwise comparisons for the 2-Hz peaks as well as for the 4-Hz peaks were significant (see Results). Since stimuli were generated pseudorandomly for each subject, and to facilitate comparisons with MEG spectra, error bars denote SEM across participants. **(D)** Bar plots of 2-Hz and 4-Hz peaks based on the MEG spectrum, showing differences between conditions. All pairwise comparisons for the 2-Hz peaks as well as for the 4-Hz peaks were significant (see Results). Error bars denote SEM across participants. **(E)** MEG channel topography of significant differences in the word-rate 2-Hz peak between conditions. Color bar denotes F statistic. The transparency mask shows significant topography clusters (p < 0.05, FWE-corrected). **(F)** MEG channel topography of significant differences in the syllable-rate 4-Hz peak between conditions. Figure legend as in (E).

The MEG spectrum (ITPC averaged across channels and conditions) also showed prominent syllable-rate (4 Hz) and word-rate (2 Hz) peaks for all three temporal conditions (isochronous, beat-based, interval-based; paired t-tests of the peaks of interest vs. neighboring frequencies; syllable-rate, all t_21_ > 2.87, all p < 0.009; word-rate: all t_21_ > 7.68, all p < 0.001; Fig. 3B). “When” predictability had a significant main effect on both the syllable-rate peaks (F_max_ = 83.57, Z_max_ = 7.82, p_FWE_ < .001) and on word-rate peaks (F_max_ = 13.29, Z_max_ = 3.98, p_FWE_ < .001; Fig. 3D). However, the syllable-rate differences showed a broad and distributed MEG topography (Fig. 3F), while the word-rate differences were more localized to left-lateralized anterior and posterior channels (Fig. 3E). Post-hoc tests revealed that, at the syllable rate, ITPC was higher in the isochronous condition than in the beat-based condition (t_21_ = 9.62, p < 0.001), and in the beat-based condition than in the interval-based condition (t_21_ = 4.16, p = 0.004). The same pattern of results was found for the word-rate ITPC (pairwise comparisons: isochronous vs. beat-based, t_21_ = 2.83, p = 0.009; beat-based vs. interval-based, t_21_ = 5.37, p < 0.001). Thus, the phase locking analysis showed a close correspondence between spectral characteristics of the stimulus envelope and the MEG responses; however, in sensor MEG data, sensitivity to word-rate peaks was relatively limited to left-lateralized channels.

### 2.3. MEG results: event-related fields and source reconstruction

To test for the temporal modulations of “what” and “when” predictions in the stimulus-evoked activity, we analyzed MEG data in the time domain and subjected ERFs to a general linear model with fixed effects “what” predictions (standard, deviant syllable, deviant word) and “when” predictions (isochronous, beat-based, interval-based). First, we found that deviants elicited higher evoked responses than standards (3-way main effect of “what” predictions, F_max_ = 12.13, Z_max_ = 4.22, p_FWE_ = 0.03). Specifically, the 3-way main effect had a nearly identical spatiotemporal distribution as the difference between deviants (pooled across deviant syllables and words) and standards (133-250 ms over left central/posterior channels; F_max_ = 24.18, Z_max_ = 4.60, p_FWE_ = 0.003; Fig. 4A), amounting to stronger deviant-evoked vs. standard-evoked ERF amplitude (T_max_ = 4.92). No significant differences were found between ERFs evoked by deviant syllables and deviant words.

**Figure 4.**
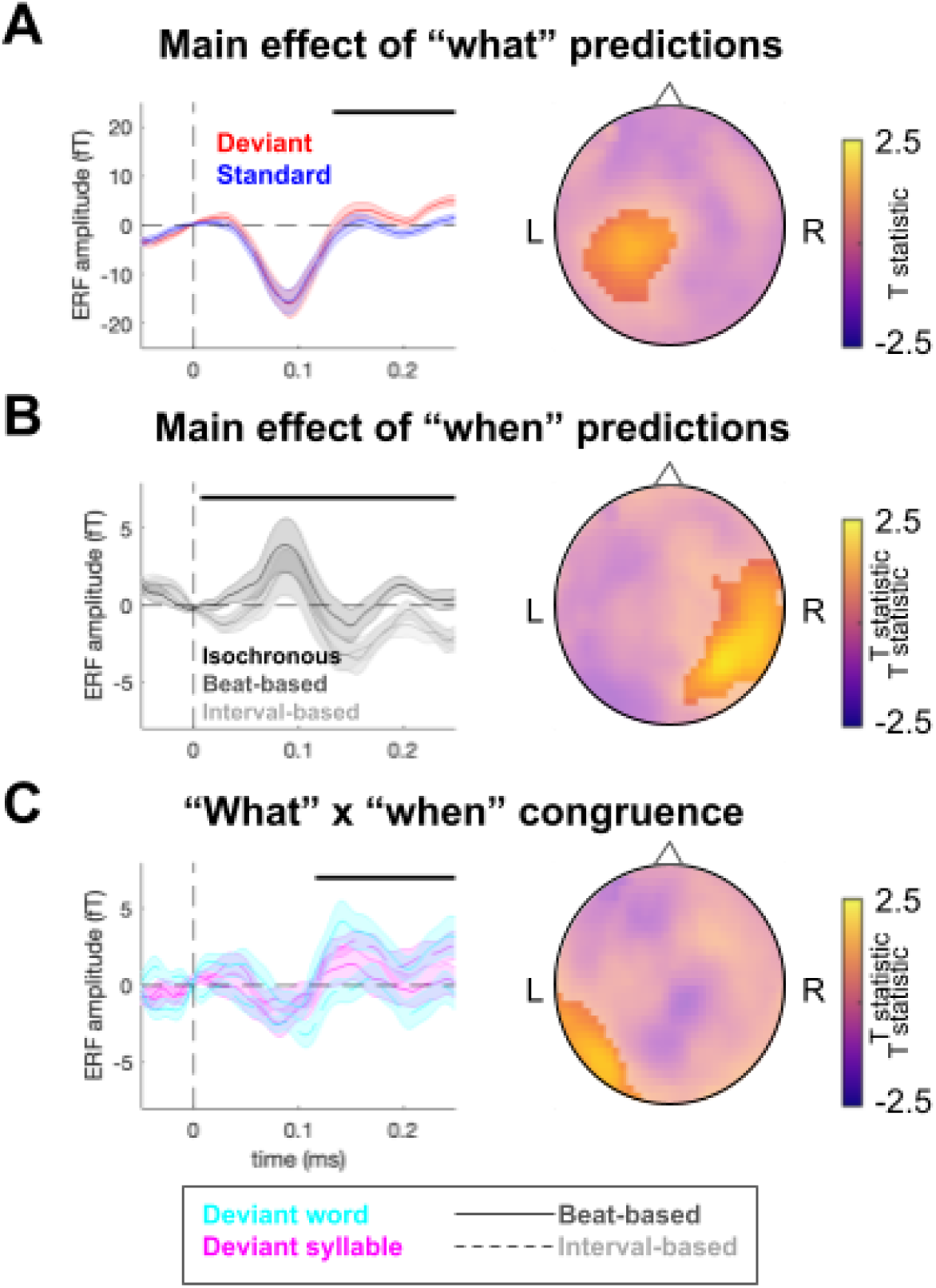
Event-related fields. **(A)** Main effect of “what”predictions (deviant vs. standard). Left panels: time courses of ERFs averaged over the spatial topography clusters shown in the right panels. Shaded area denotes SEM across participants. Black horizontal bar denotes p_FWE_ < 0.05. Right panels: spatial distribution of the main effect. Color bar: F value. **(B)** Main effect of “when” predictions (isochronous vs. interval-based vs. beat-based). Figure legend as in (A). **(C)** Congruency interaction between “what” predictions (deviant syllable vs. deviant word) and “when” predictions (interval-based vs. beat-based).

To source-localize the ERF effect found for deviants vs. standards, we reconstructed the MEG topography of evoked responses in source space and tested for differences in 3D source maps using a GLM (see Methods section 4.7 for details). Overall, averaged across stimulus types, source reconstruction could explain 88.34 ± 3.40% of sensor-space variance (mean ± SEM across participants). We identified stronger activity estimates for deviant stimuli (collapsed across deviants) vs. standard stimuli in a range of sources (Table 1; Fig. 5A), including bilateral STG, SPL and IFG, the left angular gyrus (ANG), and the right supramarginal gyrus (SMG).

**Figure 5.**
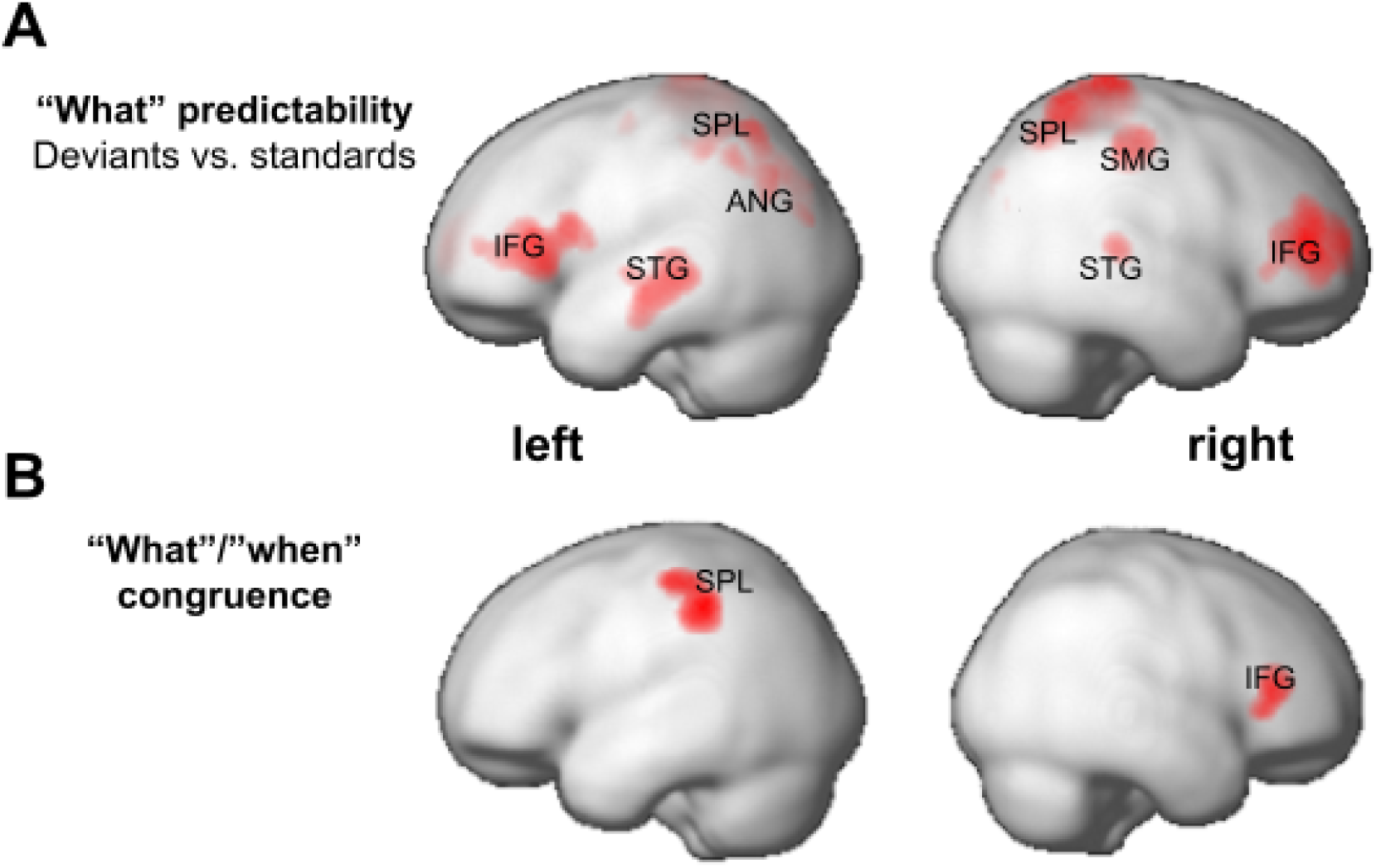
Source reconstruction. **(A)** Regions showing a significant main effect of “what” predictions (deviant vs. standard). **(B)** Regions showing a significant congruency effect between “what” predictions (deviant syllable vs. deviant word) and “when” predictions (interval-based vs. beat-based). STG: superior temporal gyrus; ANG: angular gyrus; SPL: superior parietal lobule; SMG: supramarginal gyrus; IFG: inferior frontal gyrus. Left and right hemispheres are shown in separate columns.

**Table 1.** Source reconstruction results. Summary statistics of all clusters showing significant differences between conditions (p_FWE_ < 0.05).

| Effect | Region label | Peak MNI coordinates | Voxel extent | F<sub>max</sub> | Z<sub>max</sub> | p<sub>FWE</sub> cluster-level |
| --- | --- | --- | --- | --- | --- | --- |
| “What” predictability | Right IFG/MFG | 28 50 10 | 2992 | 71.61 | 7.38 | < 0.001 |
|  | Right SPL | 22 -50 66 | 2493 | 63.97 | 7.05 | < 0.001 |
|  | Left IFG | -48 28 8 | 3513 | 58.00 | 6.76 | < 0.001 |
|  | Left SPL | -30 -64 54 | 2299 | 45.54 | 6.09 | < 0.001 |
|  | Right SMG | 46 -28 48 | 914 | 37.98 | 5.61 | 0.001 |
|  | Left ANG | -44 -70 34 | 96 | 37.81 | 5.60 | 0.001 |
|  | Right STG | 62 -32 8 | 241 | 36.67 | 5.52 | 0.008 |
|  | Left STG/MTG | -60 -18 -16 | 614 | 33.78 | 5.32 | < 0.001 |
|  | Left ANG | -42 -56 44 | 66 | 32.86 | 5.25 | 0.002 |
| “What” / “when” congruence | Left SPL | -44 -38 48 | 672 | 65.04 | 5.85 | < 0.001 |
|  | Right IFG | 42 34 14 | 281 | 55.57 | 5.56 | 0.001 |

Second, we tested for the effect of “when” predictions on ERF amplitude. While the overall 3-way difference between conditions was not significant (all p_FWE_ > 0.05), in a planned contrast we identified a significant difference between isochronous and non-isochronous (pooled over beat-based and interval-based) conditions (Fig. 4B, F_max_ = 16.92, Z_max_ = 3.84, p_FWE_ < 0.025). Here, ERF amplitudes were stronger for isochronous vs. non-isochronous conditions (T_max_ = 4.11) between 33-250 ms over right posterior channels. Source reconstruction of this ERF effect, however, did not reveal any significant source-level clusters after correcting for multiple comparisons (all p_FWE_ > 0.05).

Finally, we tested for the ERF interaction effect between “what” predictions (i.e., the type of deviant stimulus) and “when” predictions (i.e., the type of non-isochronous temporal prediction). We observed a significant interaction, such that deviant syllables and words were associated with stronger ERFs when their timing was predictable (i.e., in the interval-based and beat-based conditions, respectively), relative to when their timing was unpredictable (i.e., in the beat-based and interval-based conditions, respectively). This effect localized to left posterior channels (Fig. 4C, time extent: 127-250 ms, F_max_ = 13.10, Z_max_ = 3.36, p_FWE_ = 0.036). Post-hoc tests on the interaction effect found that it was driven primarily by deviant words presented in the beat-based vs. interval-based conditions (t_21_ = 2.899, p = 0.009), with the remaining pairwise comparisons not reaching significance after correcting for multiple comparisons (deviant syllables presented in the beat-based vs. interval-based condition: t_21_ = -1.552, p = 0.013, uncorrected; deviant syllables vs. words presented in the beat-based condition: t_21_ = -1.197, p = 0.244; deviant syllables vs. words presented in the interval-based condition: t_21_ = 2.505, p = 0.021, uncorrected).

To source-localize the interaction of “what” and “when” predictability, we compared sources of activity evoked by deviants whose timing could be predictable (deviant syllables presented in interval-based blocks and deviant words presented in beat-based blocks against deviants whose timing was unpredictable (deviant syllables presented in beat-based blocks and deviant syllables presented in interval-based blocks. This contrast revealed significant differences in two regions (Fig. 5B): the left SPL and right IFG (see Table 1 for region coordinates and statistical information). A third cluster with the most probable anatomical label being “unknown” (peak MNI [-64 -4 22], voxel extent 833, ying in the vicinity of the left postcentral/precentral gyrus)was excluded from further analysis. All sources showed weaker activity for deviant stimuli presented in temporally congruent vs. incongruent conditions (T_min_ = -8.44).

### 2.4. MEG analysis: dynamic causal modeling

To infer the connectivity patterns mediating the effects of “what” and “when” predictions on speech processing, we used dynamic causal modeling (DCM) - a Bayesian model of effective connectivity fitted to individual participants’ spatiotemporal patterns of syllable-evoked ERFs (Auksztulewicz and Friston, 2015; David et al., 2005). DCM models evoked ERFs as arising in a network of sources. Network structure is quantified by extrinsic connections (linking distinct sources) and intrinsic connections (linking distinct populations within the same source, and amounting to neural gain modulation at each source). The analysis consisted of two steps. In a first step, we created a fully interconnected model based on the 8 regions identified in the source reconstruction (see above) as well as bilateral AC (see Methods) and optimized its extrinsic connectivity using Bayesian model reduction (BMR). This procedure pruned 75% of the connections, leaving 19 significant connections out of 76 connections of the fully interconnected model in the reduced model (p < 0.001; Fig. 6A).

**Figure 6.**
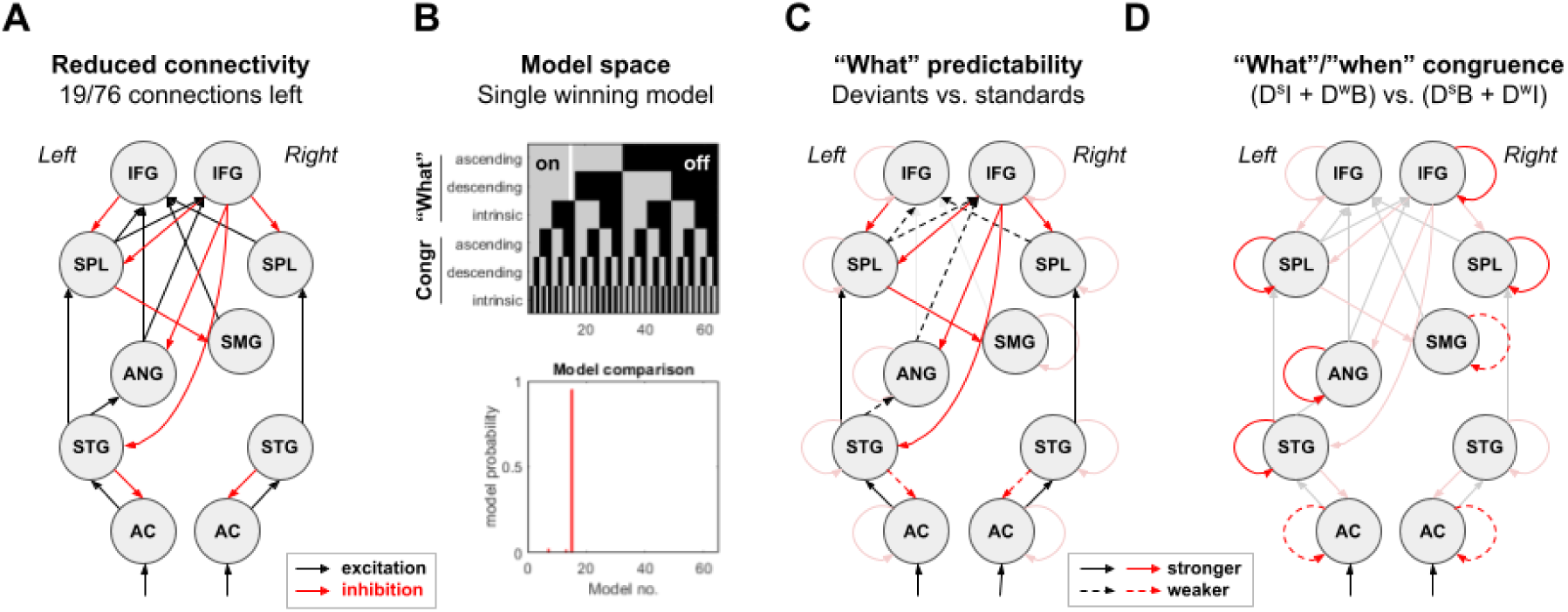
Dynamic causal modeling. **(A)** Anatomical model including the 8 regions identified in the source reconstruction analysis (bilateral superior temporal gyrus, STG; superior parietal lobule, SPL; inferior frontal gyrus, IFG; left angular gyrus, ANG; right supramarginal gyrus, SMG) as well as bilateral auditory cortex (AC). The figure depicts a model with reduced anatomical connectivity based on Bayesian model reduction, used for subsequent modeling of condition-specific effects. Black arrows: excitatory connections; red arrows: inhibitory connections. Intrinsic (self-inhibitory) connections not shown. **(B)** Upper panel: model space showing models on the horizontal axis, and groups of connections included as free parameters (grey) or switched off (black) in each model on the vertical axis. The winning model, allowing “what” predictions to modulate ascending and descending (but not intrinsic) connections, and congruence between “what” and “when” predictions to modulate intrinsic (but not ascending or descending) connections, is shown as a white column. Lower panel: Bayesian model comparison; model probability per model. **(C)** Posterior model parameters sensitive to “what” predictions. Only significant parameters (>99.9%) shown. Black: excitatory, red: inhibitory, solid lines: stronger connectivity, dashed lines: weaker connectivity for deviants vs. standards. **(D)** Posterior model parameters sensitive to congruence between “what” and “when” predictions. Self-inhibitory intrinsic connections showed region-dependent increase (solid) or decrease (dashed) for deviants temporally predicted at congruent vs. incongruent time scales.

In a second step, we took the reduced model and allowed extrinsic connections to vary systematically between “what” and “when” prediction conditions. Specifically, since in the source reconstruction results we only identified significant differences in source maps related to (1) all deviants vs. standards and (2) congruent vs. incongruent “what” and “when” predictions, we considered these two factors as possible modulatory effects of extrinsic and/or intrinsic connectivity. Bayesian model comparison of the 64 resulting models revealed a single winning model, in which “what” predictions modulated only extrinsic connections, while its congruence with “when” predictability modulated only intrinsic connections (Fig. 6B). The difference in the free-energy approximation to log-model evidence between the winning model and the next-best model (log Bayes factor) was 3.701, amounting to 97.53% probability that the winning model outperforms the next-best model.

Based on the winning model, we then inferred its significant connectivity parameters. “What” predictability significantly modulated a subset of extrinsic connections (Fig. 6C; see Table 2 for statistical information). Specifically, deviants were linked to increased ascending bilateral connectivity at multiple levels in the auditory hierarchy, from AC via STG to SPL. Conversely, ascending connectivity decreased for cross-hemispheric connections at higher levels of the hierarchy, from SPL to IFG and from the left STG via ANG to right IFG. Deviants also modulated descending connectivity, such that top-down inhibition increased at higher levels of the hierarchy (from bilateral IFG to SPL, ANG, and SMG) and decreased at lower levels of the hierarchy (from STG to AC).

**Table 2.**
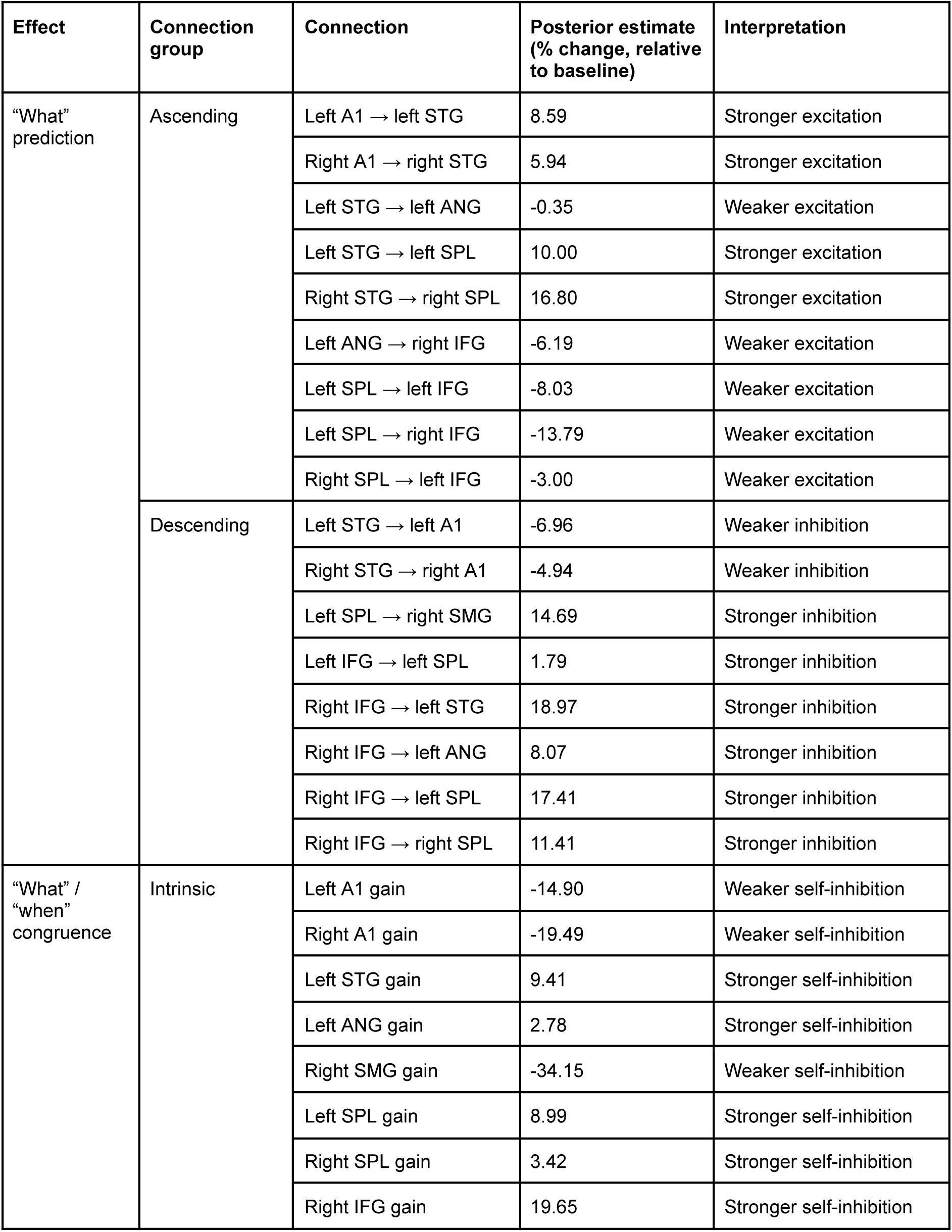
Dynamic causal modeling results. Summary of significant condition-specific effects on connectivity estimates (p < 0.001).

Finally, the congruence of “what” and “when” predictions exclusively modulated a subset of intrinsic connections (Fig. 6D; see Table 2 for statistical information). Specifically, deviant syllables and words predictable in time were linked to increased gain (decreased self-inhibition) in bilateral AC and the right SMG, and decreased gain (increased self-inhibition) in most other sources in the network with the exception of the right STG and left IFG for which no significant self-connectivity modulation was found. This model explained 81.44 ± 1.62% of the variance of spatiotemporal ERF patterns (mean ± SEM across participants).

## 3. Discussion

In the current study, we observed a contextual modulation of the neural responses to deviations from predicted speech contents, dependent on their temporal predictability. This modulation showed that faster “when” (interval-based) predictions amplified responses to deviant syllables, and slower “when” (beat-based) predictions amplified responses to deviant words. This implies a congruence effect in the processing of “what” and “when” predictions across hierarchical levels of speech processing whereby smaller processing units (syllables) are modulated at faster rates and larger processing units (words) are modulated at slower rates. However, the interactive effects between “what” and “when” predictions on evoked neural responses did not differentiate between hierarchical levels, and were instead linked to a shared network of sources including the left SPL and right IFG. In the connectivity analysis, these modulatory effects of “when” predictability on “what” mismatch responses were best explained by widespread gain modulations, including stronger sensitivity of early auditory regions bilaterally, and right SMG, as well as weaker sensitivity of most of the temporo-fronto-parietal network. Thus, our findings suggest that the interactions between “what” and “when” predictions, while contextually specific, are facilitated by a common and distributed cortical network, independent of the hierarchical level of these predictions.

Mismatch responses to unpredicted speech contents are well documented in neuroimaging studies and have been found in a range of cortical regions including the STG (Rothermich and Kotz, 2013), the SMG (Celsis et al., 1999), and - in case of nonsense words - bilateral IFG, largely matching our findings (Wilson et al., 2015). Beyond these regions, we also found sensitivity to “what” prediction violations in the angular gyrus, consistent with its role in semantic processing (Seghier, 2013), and the SPL, part of the dorsal attentional network previously linked to statistical learning of artificial speech streams (Sengupta et al., 2019). Our dynamic causal modeling of mismatch responses suggested a consistent pattern of connectivity modulations, with qualitative differences between lower and higher levels of the cortical hierarchy. At the lower levels (from A1 to STG and from STG to SPL), speech deviants were linked to stronger ascending excitation and weaker descending inhibition, consistent with increased forward prediction errors signaled to short-term violations of speech predictions by auditory regions in the temporal lobe (Caucheteux et al., 2023). Conversely, at the higher levels of the hierarchy (in the frontoparietal network), speech deviants were associated with weaker ascending connectivity and stronger descending connectivity, possibly reflecting internal attentional orienting to unpredicted speech contents (Lückmann et al., 2014; Reiche et al., 2013).

Temporal predictability of speech sounds was found to modulate behavior in an incidental task (with RTs being shortest following isochronous vs. non-isochronous sequences), consistent with previous findings. (Morillon et al., 2016). We also found that temporal predictability (in isochronous sequences) increased sound-evoked ERF amplitude (Bouwer et al., 2016), albeit to a similar extent relative to both beat-based and interval-based non-isochronous sequences. A previous EEG study comparing these two types of temporal predictions found largely comparable behavioral and neural effects, with the differences limited to sounds presented at unexpected times (but not to sounds presented at expected times) in beat-based sequences (Bouwer et al., 2020). In our study, differences between these two types of sequences were found in the analysis of frequency-domain effects of “when” predictions. Frequency-domain analyses indicated a close alignment between the MEG spectrum and the stimulus spectrum. However, while the syllable-rate effect was distributed over a large number of channels, the word-rate effect was predominantly observed over the left hemisphere, suggesting that the statistical learning of speech sequences can exert asymmetric effects. This is consistent with previous reports of left-hemispheric contributions to speech segmentation based on statistical regularities (Cunillera et al., 2009; López-Barroso et al., 2013). Similarly, studies using phase-locking measures to quantify entrainment at suprasyllabic time scales showed more pronounced differences in the left hemisphere (Ding et al., 2017; Har-Shai Yahav and Zion Golumbic, 2021), although these pertained to multi-word phrases rather than single words, and were based on familiar disyllabic words rather than recently learned pseudowords. Interestingly, the left-hemispheric dominance found for the word-rate entrainment in the current study contrasts with the right-hemispheric dominance found for tone-pair entrainment in a similar study based on non-speech musical sequences rather than on speech stimuli (Cappotto et al., 2023), suggesting differential entrainment between speech and non-speech sequences.

Besides their distinct individual effects on neural activity, predictions related to speech contents and timing demonstrated interactive effects, such that temporally predictable deviant sounds yielded stronger ERF amplitudes than temporally unpredictable deviant sounds. This interaction was specific with respect to the hierarchical level of speech organization and its respective time scale, such that deviant words had a stronger amplitude following beat-based predictions, whereas deviant syllables had a stronger amplitude following interval-based predictions. These results build upon prior research indicating that temporal predictions increase MMRs (Jalewa et al., 2021; Lumaca et al., 2019; Takegata and Morotomi, 1999; Todd et al., 2018; Yabe et al., 1997) and demonstrate that these modulatory effects align consistently with expected time points, irrespective of the specific nature of the “when” predictions (whether interval-based or beat-based). The interactive ERF effects were source-localized to two regions, the right IFG and left SPL, where deviants presented at predictable latencies were linked to lower source activity estimates than deviants presented at unpredictable latencies. Identifying these two regions as mediating interactions between “what” and “when” predictions extends the results of previous studies, where the left parietal cortex was found to be involved in integrating “what” and “when” information in speech processing (Orpella et al., 2020), while the right IFG was found to be inhibited by regular speech timing (metrical context) (Rothermich and Kotz, 2013). Our DCM results suggest that the interactive effects of “what” and “when” predictions were subserved primarily by gain modulation, including weaker gain at hierarchically higher regions, including the right IFG and left SPL as identified in the source reconstruction. This relative attenuation at hierarchically higher levels was accompanied by stronger gain at lowest levels of the network including bilateral A1, consistent with previously reported gain-amplifying effects of temporal orienting on auditory processing (Auksztulewicz et al., 2019; Auksztulewicz and Friston, 2015; Morillon et al., 2016), as well as in the SMG, previously shown to be involved in processing temporal features of speech (Geiser et al., 2008).

This MEG study analyzing speech sequences aligns closely with findings from Cappotto et al. (2023), who used similar methods to examine EEG responses to musical sequences. In both cases, faster and slower “when” predictions intensified the deviant responses to unexpected elements at the appropriate hierarchical levels—for speech, syllables vs. disyllabic words; for music, single tones vs. tone pairs. Additionally, these studies found that the interactive effects of “what” and “when” predictions are associated with the left superior parietal lobule (SPL). However, our study also implicates the right inferior frontal gyrus (IFG) as a potential source of these sensor-level effects. Furthermore, both studies reported that the sensor-level effects could be attributed to increased gain in bilateral auditory cortices and reduced gain in other network nodes. The EEG study further linked these effects to changes in forward connectivity. Collectively, these findings suggest that the interactions between “what” and “when” predictions are consistent across different stimulus domains (speech and music) and data modalities (MEG and EEG). They reveal largely overlapping cortical sources and connectivity modulations. Therefore, despite the inherent differences in temporal modulation patterns and acoustic features between music and speech (Ding et al., 2017; Siegel et al., 2012; Albouy et al., 2020; Zatorre, 2022), these common underlying mechanisms support the hypothesis that the interactions between “what” and “when” predictions are broadly generalizable across stimulus characteristics.

Taken together, our study complements recent model-based reports of cortical hierarchies aligning with speech processing hierarchies (Caucheteux et al., 2023; Schmitt et al., 2021) and suggests that while “what” and “when” predictions may jointly modulate speech processing, their interactions are not necessarily expressed at different levels of the cortical hierarchy. Instead, the effects of temporal regularities on unexpected speech sounds may be subserved by a common set of frontoparietal regions, reflecting attention-like amplification of mismatch responses due to temporal predictions (Auksztulewicz et al., 2019; Auksztulewicz and Friston, 2015) and irrespective of the contents of mispredicted stimuli. Rather than requiring dedicated resources to integrate “what” and “when” features separately at each stage of hierarchical speech processing, such a generic mechanism may help integrate streams of information across hierarchical levels.

## 4. Methods

### 4.1. Participant sample

A total of 24 participants took part in the study upon written informed consent. Two participants did not complete the study, resulting in data from 22 participants taken into analysis (13 females, 9 males; median age 28, range 21-35 years; all right-handed). The study adhered to protocols approved by the Ethics Board of the Goethe-University Frankfurt am Main. All participants confirmed normal hearing in their self-reports, and none reported any current or past neurological or psychiatric disorders.

### 4.2. Stimulus and task description

The experimental paradigm used auditory sequences which were manipulated with respect to “what” and “when” predictions at two levels each: “what” predictions of single syllables vs. disyllabic pseudowords; “when” predictions at a faster time scale of 4 Hz vs. a slower time scale of 2 Hz. This orthogonal manipulation enabled an analysis of their interactions at each level. Prior to the experimental task, participants implicitly learned 6 disyllabic pseudowords (“tupi”, “robi”, “daku”, “gola”, “zufa”, “somi”; Fig. 1A). The syllables were taken from a database of Consonant-Vowel syllables (Ives et al., 2005) and resynthesized using an open-source vocoder, STRAIGHT (Kawahara, 2006) for MATLAB R2018b (MathWorks) to match their duration (166 ms), fundamental frequency (F0 = 150 Hz), and sound intensity.

In the implicit learning task (administered outside of the MEG scanner but immediately prior to the MEG recording session), participants were exposed to continuous auditory streams of the 6 pseudowords presented in a random order. The stimulus onset asynchrony (SOA) between each two syllables was set to 250 ms, resulting in an isochronous syllable rate of 4 Hz. The stream was 120 s long, amounting to 80 occurrences of each pseudoword. Following stream offset, participants were asked to discriminate “correct” pseudowords (e.g. “tupi”) from “incorrect” pseudowords (e.g., “pitu”, “turo”, “tuku”). Each participant performed 60 trials of the pseudoword discrimination task.

Following the implicit learning task, participants were exposed to the main experimental paradigm in the MEG scanner. They were asked to listen to continuous sequences of 4 unique pseudowords (e.g., “tupirobidakugola”). As a cover task, to monitor participants’ attention, we asked them to detect immediate syllable repetitions (e.g., “tupirobidakugogo”), present in 6.6% sequences, immediately after repetition onset. These trials were rejected from subsequent analysis of neural data.

The sequences presented in the MEG scanner were independently manipulated with respect to “what” and “when” predictions. “What” prediction manipulations had three levels (Fig. 1B): (1) in standard sequences (66.6%), only correct pseudowords were used (e.g., “tupirobidakugola”); (2) in “deviant syllable” sequences (13.3%), the final syllable of the sequence was replaced with a syllable belonging to a different pseudoword (e.g., “tupirobidakugo*fa*”), such that upon hearing the syllable “go”, the prediction of the subsequent syllable “la” is violated by the syllable “fa”; (3) in “deviant word” sequences (13.3%), the penultimate syllable of the sequence was replaced with a syllable which should be a final syllable of a different pseudoword (e.g., “tupirobidaku*fa*la”), creating an unpredictable disyllabic pseudoword (“fala”). “What” predictions were manipulated at a sequence-by-sequence level, such that different types of sequences were presented in a random order. Each trial contained 7 seamlessly concatenated sequences, amounting to 14 s per trial. Trials were separated by an inter-trial interval of 1 s. To match the positions and timing of deviants and standards in the sequences, for each deviant syllable/word a designated standard syllable/word was drawn from the same position (i.e., penultimate or final syllable) of a standard sequence with matched timing.

Independently, “when” prediction manipulations also had three levels (Fig. 1C): (1) in isochronous sequences (33% of the trials), the SOA between consecutive syllables was fixed at 250 ms; (2) in “beat-based” sequences (33% of the trials), the SOA between pseudoword-initial syllables was fixed at 500 ms, but the SOA between pseudoword-initial and pseudoword-final syllables was jittered between 167 and 333 ms, resulting in unpredictable timing of final syllables of each pseudoword (i.e., at a faster time scale, corresponding to syllable rate) but predictable timing of pseudoword onsets (i.e., at a slower time scale, corresponding to word rate); (3) in “interval-based” sequences (33% of the trials), the SOA between pseudoword-initial and pseudoword-final syllables was fixed at 250 ms, but the SOA between pseudoword-initial syllables was jittered between 167 and 333 ms, resulting in unpredictable timing of pseudoword onsets (i.e., at the slower time scale) but predictable timing of final syllables of each pseudoword (i.e., at the faster time scale). “When” predictions were manipulated at a block level (20 trials per block). Each “when” condition was administered in 4 blocks, resulting in 12 blocks in total. Blocks were presented in a pseudorandom order, such that no immediate repetitions of the same “when” condition was allowed.

To ensure that the MEG analysis was not confounded by differences in baseline duration between temporal conditions (e.g., syllable deviants preceded by shorter/longer SOAs in the beat-based condition than in the remaining conditions), the SOAs preceding all deviant syllables and the designated standard syllables were replaced by a fixed 250 ms SOA. Therefore, the temporal predictability manipulation was limited to syllables surrounding the analyzed syllables but not to the immediately preceding syllables. The global deviant probability equaled 4.16% (including repetitions) or 3.32% (excluding repetitions) of all syllables, amounting to 80 deviant syllables per combination of “what” prediction violation (deviant word vs. syllable) and “when” prediction condition (isochronous, beat-based, interval-based).

### 4.3. Behavioral analysis

Behavioral analysis focused on accuracy and response time (RT) data derived from participant responses in the syllable repetition detection task. Single-trial RTs exceeding each participant’s median + 3 standard deviations were removed from the analysis. The remaining RTs, derived exclusively from correct trials, underwent a log transformation to achieve an approximately normal distribution, and subsequently averaged. To test for the effect of “when” predictions on behavioral performance in the repetition detection task, accuracy and mean log RTs were separately subjected to repeated-measures ANOVAs, incorporating the within-subjects factor of Time (isochronous, interval-based, beat-based). Since the behavioral task was based on repetitions but not on deviant words or syllables, “what” predictions were not included as a factor in these analyses. Post-hoc comparisons were executed through paired t-tests in MATLAB, and corrections were applied for multiple comparisons — specifically, three for accuracy and three for RTs — using a false discovery rate of 0.05.

### 4.4. MEG data acquisition and pre-processing

Participants were seated in a 275-channel whole-head CTF MEG system with axial gradients (Omega 2005, VSM MedTech Ltd., BC, Canada). The data were acquired at a sampling rate of 1200 Hz with synthetic third-order gradient noise reduction (Vrba and Robinson, 2001). For monitoring eye blinks and heart rate, four electrooculogram (EOG) and two electrocardiogram (ECG) electrodes were placed on the participant’s face and clavicles. Continuous head localization was recorded throughout the session.

Auditory stimuli were generated by an external sound card (RME) and transmitted into the MEG chamber through sound-conducting tubes which were linked to plastic ear molds (Promolds, Doc’s Proplugs). The sound pressure level was adjusted to approximately 70 dB SPL. Visual stimuli were presented using a PROPIX projector (VPixx ProPixx) and back-projected onto a semi-transparent screen positioned 60 cm from the participant’s head. Participants responded to stimuli by operating a MEG-compatible button response box (Cambridge Research Systems, LTD) with their right hand. Short breaks were administered between runs.

The continuous MEG recordings were high-pass filtered at 0.1 Hz and notch filtered between 48 Hz and 52 Hz, down-sampled to 300 Hz, and further subjected to a low-pass filter at 90 Hz (including antialiasing). All filters were 5th order zero-phase Butterworth and implemented in the SPM12 toolbox for Matlab. Based on continuous head position measurement inside the MEG scanner, we calculated 6 movement parameters (3 translations and 3 rotations) (Stolk et al., 2013), which were regressed out from each MEG channel using linear regression. Eyeblink artifacts were automatically detected based on the vertical EOG, and removed by subtracting the two top spatiotemporal principal components of eyeblink-evoked responses from all MEG channels (Ille et al., 2002). Heartbeat artifacts were automatically detected based on ECG and removed in the same manner. The cleaned signals were subsequently subjected to separate analyses in the frequency domain (phase-locking) and the time domain (event-related fields).

### 4.5. MEG analysis: phase locking

To investigate whether speech sequences exhibit distinct spectral peaks in neural responses at both the syllable rate (4 Hz) and the word rate (2 Hz), we conducted a frequency domain analysis. The continuous data were segmented into epochs spanning from the onset to the offset of each trial (speech sequence). For each participant, channel, and sequence, we computed the Fourier spectrum of MEG signals recorded during that specific sequence. Following previous research (Ding and Simon, 2013), to assess phase consistency within each condition we computed the inter-trial phase coherence (ITPC) for each temporal condition (isochronous, interval-based, beat-based) using the following equation:

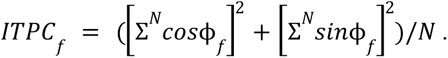

In the equation above, Φ_f_ denotes the Fourier phase for a given frequency f, and N denotes the number of trials (here: sequences; 80 per condition).

We applied ITPC in the aperiodic conditions as well, based on previous findings that temporally consistent slow ramping neural activity (such as the contingent negative variation) can produce significant ITPC values even in aperiodic (interval-based) sequences, as demonstrated in previous research (Breska and Ivry, 2018). We also assessed the neural phase consistency in response to both periodic and aperiodic stimuli conditions by calculating the ITPC based on the raw stimulus waveform.

In the initial analysis, ITPC estimates were averaged across MEG channels. To assess the presence of statistically significant spectral peaks, ITPC values at the syllable rate (4 Hz) and word rate (2 Hz) were compared against the mean of ITPC values at their respective neighboring frequencies (syllable rate: 3.93 and 4.07 Hz; word rate: 1.929 and 2.071 Hz) using paired t-tests.

Additionally, to examine whether spectral peaks at the syllable rate and word rate observed at individual MEG channels exhibited modulations due to temporal predictability, spatial topography maps of single-channel ITPC estimates were transformed into 2D images. These images were then smoothed with a 5 x 5 mm full-width-at-half-maximum (FWHM) Gaussian kernel to match the expected spatial scale of MEG sensor-level data. The smoothed images were subjected to repeated-measures ANOVAs, separately for syllable-rate and word-rate ITPC estimates, incorporating a within-subjects factor of Time (isochronous, interval-based, beat-based). This analysis was implemented in SPM12 as a general linear model (GLM). To address multiple comparisons and ITPC correlations across neighboring channels, statistical parametric maps were thresholded at p < 0.005 and corrected for multiple comparisons over space at a cluster-level p_FWE_ < 0.05, following random field theory assumptions (Kilner et al., 2005). Repeated-measures parametric tests were selected based on previous literature using ITPC (Sokoliuk et al., 2021), assuming that differences in ITPC values between conditions follow a normal distribution. Post-hoc tests were conducted at a Bonferroni-corrected FWE threshold (0.05 / 3 pairwise comparisons per rate).

### 4.6. MEG analysis: event-related fields

For the analysis in the time domain, the data underwent segmentation into epochs spanning from -50 ms before to 250 ms after the onset of deviant/standard syllables. To prevent contamination from the temporally structured presentation, baseline correction was applied from -25 ms to 25 ms, following a recently published approach (Fitzgerald et al., 2021). The data were then denoised using the “Dynamic Separation of Sources” algorithm (de Cheveigné and Simon, 2008), aimed at minimizing the impact of noisy channels. Condition-specific event-related fields (ERFs) corresponding to syllable and word deviants and the respective standards, presented in each of the three temporal conditions, were computed using robust averaging across trials to mitigate the influence of outlier trials. This analysis was conducted with the SPM12 toolbox and included a low-pass filter at 48 Hz (5th order zero-phase Butterworth) to derive the final ERFs. The ERFs were then subjected to univariate analysis to assess the effects of “what” and “when” predictions on evoked responses.

The ERF data were transformed into 3D images (2D for spatial topography; 1D for time) which underwent spatial smoothing using a 5 x 5 mm FWHM Gaussian kernel. Subsequently, the smoothed images were entered into a GLM, which implemented a 3 x 3 repeated-measures ANOVA with within-subject factors Content (standard, deviant syllable, deviant word) and Time (isochronous, interval-based, beat-based). In addition to testing for the two main effects and a general 3 x 3 interaction, we also tested for the following planned contrasts: (1) deviant vs. standard, (2) isochronous vs. non-isochronous, (3) an interaction contrast isolating the congruence effect. This last contrast aimed to investigate whether “when” predictions specifically influenced the amplitude of mismatch responses evoked by deviants presented at a time scale congruent with “when” predictions — i.e., deviant syllables in the interval-based condition and deviant words in the beat-based conditions. This involved testing for a 2 x 2 interaction between Content (deviant syllable, deviant word) and Time (interval-based, beat-based).

To address multiple comparisons and ERF amplitude correlations across neighboring channels and time points, statistical parametric maps were thresholded at p < 0.005 and corrected for multiple comparisons over space and time at a cluster-level p_FWE_ < 0.05, following random field theory assumptions (Kilner et al., 2005).

### 4.7. MEG analysis: source reconstruction

Source reconstruction was conducted under group constraints (Litvak and Friston, 2008), enabling the estimation of source activity at the individual participant level by assuming that activity is reconstructed within the same subset of sources for each participant, thereby reducing the impact of outliers. An empirical Bayesian beamformer (Belardinelli et al., 2012; Little et al., 2018; Wipf and Nagarajan, 2009) was employed for estimating sources based on the entire post-stimulus time window (0-250 ms). Given the principal findings identified in the ERF analysis—specifically, a difference between ERFs elicited by deviants and standards; a difference between ERFs elicited in isochronous and non-isochronous contexts; and an interaction between deviant type and temporal condition—we focused on comparing source estimates related to these effects.

For the analysis of the difference between deviants and standards, as well as the difference between stimuli presented in isochronous and non-isochronous sequences, source estimates were extracted for the 33-250 ms time window (based on the results of the ERF analysis; see below). These estimates were converted into 3D images with three spatial dimensions and then smoothed using a 5 x 5 x 5 mm Full-Width-at-Half-Maximum (FWHM) Gaussian kernel. The smoothed images were entered into a GLM that implemented a 3 x 3 repeated-measures ANOVA with within-subjects factors of Content (standard, deviant syllable, deviant word) and Time (isochronous, interval-based, beat-based). Parametric tests based on a GLM were employed as an established method for analyzing MEG source reconstruction maps (Litvak et al., 2011).

For the analysis of the interaction between deviant type and temporal condition, source estimates were extracted for the 127-250 ms time window (based on the results of the ERF analysis) and processed as described above. Smoothed images were then entered into a GLM implementing a 2 x 2 repeated-measures ANOVA with within-subjects factors of Content (deviant syllable, deviant word) and Time (interval-based, beat-based). To address multiple comparisons and source estimate correlations across neighboring voxels, statistical parametric maps were thresholded and corrected for multiple comparisons over space at a cluster-level p_FWE_ < 0.005 (minimum voxel extent: 64 voxels), adhering to random field theory assumptions (Kilner et al., 2005). Source labels were assigned using the Neuromorphometrics probabilistic atlas, implemented in SPM12.

### 4.8. MEG analysis: dynamic causal modeling

Dynamic causal modeling (DCM) was employed to estimate connectivity parameters at the source level, specifically related to the general processing of mismatch responses (deviant vs. standard) and the contextual interaction between “what” and “when” predictions (deviant syllable in the interval-based condition and deviant word in the beat-based condition vs. deviant syllable in the beat-based condition and deviant word in the interval-based condition). Since no sources were identified in the source reconstruction of the main effect of “when” predictions (see below), this effect was not included in the model. DCM, a form of effective connectivity analysis, utilizes a generative model to map sensor-level data (in this case, ERF time series across MEG channels) to source-level activity (David et al., 2005). The generative model encompasses several sources representing distinct cortical regions, forming a sparsely interconnected network.

Each source’s activity is explained by neural populations based on a canonical microcircuit (Bastos et al., 2012), modeled using coupled differential equations describing changes in postsynaptic voltage and current in each population. In our study, the microcircuit consisted of four populations (superficial and deep pyramidal cells, spiny stellate cells, and inhibitory interneurons), each with a unique connectivity profile, including ascending, descending, and lateral extrinsic connectivity (connecting different sources) and intrinsic connectivity (connecting different populations within each source). The canonical microcircuit’s form and connectivity profile followed procedures established in the previous literature on the subject (Auksztulewicz et al., 2018; Auksztulewicz and Friston, 2015; Fitzgerald et al., 2021; Rosch et al., 2019; Todorovic and Auksztulewicz, 2021).

Crucially, a subset of intrinsic connections represented self-connectivity parameters, describing the neural gain of each region. Both extrinsic connectivity and gain parameters were allowed to undergo condition-specific changes to model differences between experimental conditions (deviants vs. standards and the interaction between “what” and “when” predictions). The canonical microcircuit models prior connection weights for all ascending and lateral connections as excitatory, and for descending and intrinsic connections as inhibitory, based on the previous literature linking descending connections to predictive suppression, and intrinsic connections to self-inhibitory gain control (Bastos et al., 2012). However, these priors can be overridden by the data likelihood, possibly resulting in ascending/lateral inhibition and descending/intrinsic excitation if this maximizes model evidence.

In this study, we employed DCM to fit the individual participants’ ERFs specific to each condition within the 0-250 ms timeframe. Drawing on the results of source reconstruction (refer to the Results section) and prior research (Garrido et al., 2008), we integrated 10 sources into the cortical network. These 10 sources included 8 regions identified in the source reconstruction (Fig. 5) based on their peak MNI coordinates: bilateral superior temporal gyrus (STG; left, [-60 -18 -16]; right, [62 -32 8]), left angular gyrus ([-44 -70 34], right supramarginal gyrus ([44 -26 48]), bilateral superior parietal lobule (left, [-30 -64 54]; right, [22 -50 66]), and bilateral inferior frontal gyrus (left, [-48 28 8]; right, [42 34 14]). Additionally, we included 2 regions corresponding to bilateral primary auditory cortex for anatomical plausibility (Garrido et al., 2008) (A1; MNI coordinates: left, [-56 -12 -2]; right, [60 -14 18]). The A1 coordinates were based on local maxima in the source reconstruction contrast maps for which probabilistic labeling returned “planum polare” (left A1 coordinates) or “planum temporale” (right A1 coordinates). To evaluate model fits, we utilized the free-energy approximation to model evidence, penalized by model complexity.

The analysis followed a sequential approach: initially, model parameters encompassing only extrinsic connections were estimated based on all experimental conditions, without modeling differences between conditions. The aim of this initial step was to find the optimal connectivity pattern between sources, providing the best fit of the data. In a second step, condition-specific changes in both extrinsic and intrinsic connections were optimized at the individual participant level. In both steps, models were fitted to individual participants’ data; significant parameters (connection weights) were inferred at the group level using parametric empirical Bayes (PEB) and models were optimized using Bayesian model reduction (BMR) (Friston et al., 2016), as described below.

At the individual participant level, models were fitted to ERF data considering two factors: “what” predictions (all deviants vs. standards) and the contextual interaction between “what” and “when” predictions (deviant syllable in the interval-based condition and deviant word in the beat-based condition vs. deviant syllable in the beat-based condition and deviant word in the interval-based condition). At this stage, all extrinsic and intrinsic connections were included in the network, representing a “full” model. Due to the potential for local maxima in model inversion within DCM, the group level analysis implemented PEB. This involved inferring group-level parameters by (re)fitting the same “full” models to individual participants’ data. The assumption underlying this step was that model parameters should be normally distributed in the participant sample, helping to mitigate the impact of outlier participants. For model comparison, BMR was applied, contrasting the “full” models against a range of “reduced” models where certain parameters were fixed to zero. This led to the creation of a model space encompassing different combinations of parameters.

In the first step (optimizing extrinsic connections independent of conditions), we used BMR to prune the extrinsic connectivity matrix. The free-energy approximation to log-model evidence was employed to score each model with a given extrinsic connectivity parameter set to 0, relative to the full model. This approach resulted in Bayesian confidence intervals for each parameter, indicating the uncertainty of parameter estimates. Parameters with 99.9% confidence intervals spanning either side of zero (equivalent to p < 0.001) were deemed statistically significant.

In the second step (optimizing the modulation of extrinsic and intrinsic connections by experimental conditions), a total of 64 models were generated. The model space was designed in a factorial manner, such that the following six groups of connections were set as free parameters or fixed to zero independent of each other: (1) ascending connectivity modulation by “what” predictions; (2) descending connectivity modulation by “what” predictions; (3) intrinsic connectivity modulation by “what” predictions; (4) ascending connectivity modulation by “what” and “when” congruence; (5) descending connectivity modulation by “what” and “when” congruence; (6) intrinsic connectivity modulation by “what” and “when” congruence. The resulting 64 (2^6) models were fitted using BMR (switching off subsets of parameters of the full model) and compared using Bayesian model selection. Since a single model was identified as winning (see Results), its posterior parameters were inspected. Parameters with 99.9% non-zero confidence intervals were treated as statistically significant.

## Acknowledgments

This work has been supported by the European Commission’s Marie Skłodowska-Curie Global Fellowship (750459 to R.A.); a grant from the European Commission / Hong Kong Research Grants Council Joint Research Scheme (9051402 to R.A. and J.S.); a grant from the German Science Foundation (AU 423/2-1 to R.A.).

## References

Arnal LH, Giraud A-L. 2012. Cortical oscillations and sensory predictions. Trends Cogn Sci 16:390–398.

Auksztulewicz R, Friston K. 2015. Attentional Enhancement of Auditory Mismatch Responses: a DCM/MEG Study. Cereb Cortex 25:4273–4283.

Auksztulewicz R, Myers NE, Schnupp JW, Nobre AC. 2019. Rhythmic Temporal Expectation Boosts Neural Activity by Increasing Neural Gain. J Neurosci 39:9806–9817.

Auksztulewicz R, Schwiedrzik CM, Thesen T, Doyle W, Devinsky O, Nobre AC, Schroeder CE, Friston KJ, Melloni L. 2018. Not All Predictions Are Equal: “What” and “When” Predictions Modulate Activity in Auditory Cortex through Different Mechanisms. J Neurosci 38:8680–8693.

Bastos AM, Usrey WM, Adams RA, Mangun GR, Fries P, Friston KJ. 2012. Canonical microcircuits for predictive coding. Neuron 76:695–711.

Belardinelli P, Ortiz E, Barnes G, Noppeney U, Preissl H. 2012. Source reconstruction accuracy of MEG and EEG Bayesian inversion approaches. PLoS One 7:e51985.

Blank H, Spangenberg M, Davis MH. 2018. Neural Prediction Errors Distinguish Perception and Misperception of Speech. J Neurosci 38:6076–6089.

Bouwer FL, Honing H, Slagter HA. 2020. Beat-based and Memory-based Temporal Expectations in Rhythm: Similar Perceptual Effects, Different Underlying Mechanisms. J Cogn Neurosci 32:1221–1241.

Bouwer FL, Werner CM, Knetemann M, Honing H. 2016. Disentangling beat perception from sequential learning and examining the influence of attention and musical abilities on ERP responses to rhythm. Neuropsychologia 85:80–90.

Breska A, Ivry RB. 2018. Double dissociation of single-interval and rhythmic temporal prediction in cerebellar degeneration and Parkinson’s disease. Proc Natl Acad Sci U S A 115:12283–12288.

Cappotto D, Luo D, Lai HW, Peng F, Melloni L, Schnupp JWH, Auksztulewicz R. 2023. “What” and “when” predictions modulate auditory processing in a mutually congruent manner. Front Neurosci 17:1180066.

Caucheteux C, Gramfort A, King J-R. 2023. Evidence of a predictive coding hierarchy in the human brain listening to speech. Nat Hum Behav 7:430–441.

Celsis P, Boulanouar K, Doyon B, Ranjeva JP, Berry I, Nespoulous JL, Chollet F. 1999. Differential fMRI responses in the left posterior superior temporal gyrus and left supramarginal gyrus to habituation and change detection in syllables and tones. Neuroimage 9:135–144.

Cope TE, Sohoglu E, Sedley W, Patterson K, Jones PS, Wiggins J, Dawson C, Grube M, Carlyon RP, Griffiths TD, Davis MH, Rowe JB. 2017. Evidence for causal top-down frontal contributions to predictive processes in speech perception. Nat Commun 8:1–16.

Cunillera T, Càmara E, Toro JM, Marco-Pallares J, Sebastián-Galles N, Ortiz H, Pujol J, Rodríguez-Fornells A. 2009. Time course and functional neuroanatomy of speech segmentation in adults. Neuroimage 48:541–553.

David O, Harrison L, Friston KJ. 2005. Modelling event-related responses in the brain. Neuroimage 25:756–770.

de Cheveigné A, Simon JZ. 2008. Denoising based on spatial filtering. J Neurosci Methods 171:331–339.

Ding N, Melloni L, Zhang H, Tian X, Poeppel D. 2016. Cortical tracking of hierarchical linguistic structures in connected speech. Nat Neurosci 19:158–164.

Ding N, Patel AD, Chen L, Butler H, Luo C, Poeppel D. 2017. Temporal modulations in speech and music. Neurosci Biobehav Rev 81:181–187.

Ding N, Simon JZ. 2013. Power and phase properties of oscillatory neural responses in the presence of background activity. J Comput Neurosci 34:337–343.

Doelling KB, Assaneo MF, Bevilacqua D, Pesaran B, Poeppel D. 2019. An oscillator model better predicts cortical entrainment to music. Proc Natl Acad Sci U S A 116:10113–10121.

Donhauser PW, Baillet S. 2020. Two Distinct Neural Timescales for Predictive Speech Processing. Neuron 105:385–393.e9.

Federmeier KD. 2007. Thinking ahead: The role and roots of prediction in language comprehension. Psychophysiology 44:491–505.

Fitzgerald K, Auksztulewicz R, Provost A, Paton B, Howard Z, Todd J. 2021. Hierarchical Learning of Statistical Regularities over Multiple Timescales of Sound Sequence Processing: A Dynamic Causal Modeling Study. J Cogn Neurosci 33:1549–1562.

Friston KJ, Litvak V, Oswal A, Razi A, Stephan KE, van Wijk BCM, Ziegler G, Zeidman P. 2016. Bayesian model reduction and empirical Bayes for group (DCM) studies. Neuroimage 128:413–431.

Garrido MI, Friston KJ, Kiebel SJ, Stephan KE, Baldeweg T, Kilner JM. 2008. The functional anatomy of the MMN: a DCM study of the roving paradigm. Neuroimage 42:936–944.

Geiser E, Zaehle T, Jancke L, Meyer M. 2008. The neural correlate of speech rhythm as evidenced by metrical speech processing. J Cogn Neurosci 20:541–552.

Gómez Varela I, Orpella J, Poeppel D, Ripolles P, Assaneo MF. 2024. Syllabic rhythm and prior linguistic knowledge interact with individual differences to modulate phonological statistical learning. Cognition 245:105737.

Hakonen M, May PJC, Jääskeläinen IP, Jokinen E, Sams M, Tiitinen H. 2017. Predictive processing increases intelligibility of acoustically distorted speech: Behavioral and neural correlates. Brain Behav 7:e00789.

Har-Shai Yahav P, Zion Golumbic E. 2021. Linguistic processing of task-irrelevant speech at a cocktail party. Elife 10. doi:10.7554/eLife.65096

Heilbron M, Armeni K, Schoffelen J-M, Hagoort P, de Lange FP. 2022. A hierarchy of linguistic predictions during natural language comprehension. Proc Natl Acad Sci U S A 119:e2201968119.

Hickok G. 2012. The cortical organization of speech processing: feedback control and predictive coding the context of a dual-stream model. J Commun Disord 45:393–402.

Hovsepyan S, Olasagasti I, Giraud A-L. 2020. Combining predictive coding and neural oscillations enables online syllable recognition in natural speech. Nat Commun 11:3117.

Ille N, Berg P, Scherg M. 2002. Artifact correction of the ongoing EEG using spatial filters based on artifact and brain signal topographies. J Clin Neurophysiol 19:113–124.

Ives DT, Smith DRR, Patterson RD. 2005. Discrimination of speaker size from syllable phrases. J Acoust Soc Am 118:3816–3822.

Jalewa J, Todd J, Michie PT, Hodgson DM, Harms L. 2021. Do rat auditory event related potentials exhibit human mismatch negativity attributes related to predictive coding? Hear Res 399:107992.

Kawahara H. 2006. STRAIGHT, exploitation of the other aspect of VOCODER: Perceptually isomorphic decomposition of speech sounds. Acoust Sci Technol 27:349–353.

Keitel A, Ince RAA, Gross J, Kayser C. 2017. Auditory cortical delta-entrainment interacts with oscillatory power in multiple fronto-parietal networks. Neuroimage 147:32–42.

Kilner JM, Kiebel SJ, Friston KJ. 2005. Applications of random field theory to electrophysiology. Neurosci Lett 374:174–178.

Klimovich-Gray A, Barrena A, Agirre E, Molinaro N. 2021. One Way or Another: Cortical Language Areas Flexibly Adapt Processing Strategies to Perceptual And Contextual Properties of Speech. Cereb Cortex 31:4092–4103.

Kösem A, Bosker HR, Takashima A, Meyer A, Jensen O, Hagoort P. 2018. Neural Entrainment Determines the Words We Hear. Curr Biol 28:2867–2875.e3.

Ling X, Sun P, Zhao L, Jiang S, Lu Y, Cheng X, Guo X, Zhu X, Zheng L. 2022. Neural Basis of the Implicit Learning of Complex Artificial Grammar with Nonadjacent Dependencies. J Cogn Neurosci 34:2375–2389.

Little S, Bonaiuto J, Meyer SS, Lopez J, Bestmann S, Barnes G. 2018. Quantifying the performance of MEG source reconstruction using resting state data. Neuroimage 181:453–460.

Litvak V, Friston K. 2008. Electromagnetic source reconstruction for group studies. Neuroimage 42:1490–1498.

Litvak V, Mattout J, Kiebel S, Phillips C, Henson R, Kilner J, Barnes G, Oostenveld R, Daunizeau J, Flandin G, Penny W, Friston K. 2011. EEG and MEG data analysis in SPM8. Comput Intell Neurosci 2011. doi:10.1155/2011/852961

López-Barroso D, Catani M, Ripollés P, Dell’Acqua F, Rodríguez-Fornells A, de Diego-Balaguer R. 2013. Word learning is mediated by the left arcuate fasciculus. Proc Natl Acad Sci U S A 110:13168–13173.

Lückmann HC, Jacobs HIL, Sack AT. 2014. The cross-functional role of frontoparietal regions in cognition: internal attention as the overarching mechanism. Prog Neurobiol 116:66–86.

Lumaca M, Trusbak Haumann N, Brattico E, Grube M, Vuust P. 2019. Weighting of neural prediction error by rhythmic complexity: A predictive coding account using mismatch negativity. Eur J Neurosci 49:1597–1609.

Mai G, Wang WS-Y. 2023. Distinct roles of delta- and theta-band neural tracking for sharpening and predictive coding of multi-level speech features during spoken language processing. Hum Brain Mapp 44:6149–6172.

Morillon B, Schroeder CE, Wyart V, Arnal LH. 2016. Temporal Prediction in lieu of Periodic Stimulation. J Neurosci 36:2342–2347.

Obleser J, Kayser C. 2019. Neural Entrainment and Attentional Selection in the Listening Brain. Trends Cogn Sci 23:913–926.

Oganian Y, Chang EF. 2019. A speech envelope landmark for syllable encoding in human superior temporal gyrus. Sci Adv 5:eaay6279.

Oganian Y, Kojima K, Breska A, Cai C, Findlay A, Chang E, Nagarajan SS. 2023. Phase Alignment of Low-Frequency Neural Activity to the Amplitude Envelope of Speech Reflects Evoked Responses to Acoustic Edges, Not Oscillatory Entrainment. J Neurosci 43:3909–3921.

Orpella J, Ripollés P, Ruzzoli M, Amengual JL, Callejas A, Martinez-Alvarez A, Soto-Faraco S, de Diego-Balaguer R. 2020. Integrating when and what information in the left parietal lobe allows language rule generalization. PLoS Biol 18:e3000895.

Petersson K-M, Folia V, Hagoort P. 2012. What artificial grammar learning reveals about the neurobiology of syntax. Brain Lang 120:83–95.

Poeppel D, Assaneo MF. 2020. Speech rhythms and their neural foundations. Nat Rev Neurosci 21:322–334.

Reiche M, Hartwigsen G, Widmann A, Saur D, Schröger E, Bendixen A. 2013. Involuntary attentional capture by speech and non-speech deviations: a combined behavioral-event-related potential study. Brain Res 1490:153–160.

Riecke L, Formisano E, Sorger B, Başkent D, Gaudrain E. 2018. Neural Entrainment to Speech Modulates Speech Intelligibility. Curr Biol 28:161–169.e5.

Rimmele JM, Sun Y, Michalareas G, Ghitza O, Poeppel D. 2023. Dynamics of Functional Networks for Syllable and Word-Level Processing. Neurobiol Lang (Camb) 4:120–144.

Rosch RE, Auksztulewicz R, Leung PD, Friston KJ, Baldeweg T. 2019. Selective Prefrontal Disinhibition in a Roving Auditory Oddball Paradigm Under N-Methyl-D-Aspartate Receptor Blockade. Biol Psychiatry Cogn Neurosci Neuroimaging 4:140–150.

Rothermich K, Kotz SA. 2013. Predictions in speech comprehension: fMRI evidence on the meter-semantic interface. Neuroimage 70:89–100.

Ryskin R, Nieuwland MS. 2023. Prediction during language comprehension: what is next? Trends Cogn Sci 27:1032–1052.

Schmitt L-M, Erb J, Tune S, Rysop AU, Hartwigsen G, Obleser J. 2021. Predicting speech from a cortical hierarchy of event-based time scales. Science Advances. doi:10.1126/sciadv.abi6070

Seghier ML. 2013. The angular gyrus: multiple functions and multiple subdivisions. Neuroscientist 19:43–61.

Sengupta P, Burgaleta M, Zamora-López G, Basora A, Sanjuán A, Deco G, Sebastian-Galles N. 2019. Traces of statistical learning in the brain’s functional connectivity after artificial language exposure. Neuropsychologia 124:246–253.

Shain C, Blank IA, van Schijndel M, Schuler W, Fedorenko E. 2020. fMRI reveals language-specific predictive coding during naturalistic sentence comprehension. Neuropsychologia 138:107307.

Sohoglu E, Davis MH. 2016. Perceptual learning of degraded speech by minimizing prediction error. Proc Natl Acad Sci U S A 113:E1747–56.

Sokoliuk R, Degano G, Banellis L, Melloni L, Hayton T, Sturman S, Veenith T, Yakoub KM, Belli A, Noppeney U, Cruse D. 2021. Covert Speech Comprehension Predicts Recovery From Acute Unresponsive States. Ann Neurol 89:646–656.

Stolk A, Todorovic A, Schoffelen J-M, Oostenveld R. 2013. Online and offline tools for head movement compensation in MEG. Neuroimage 68:39–48.

Su Y, MacGregor LJ, Olasagasti I, Giraud A-L. 2023. A deep hierarchy of predictions enables online meaning extraction in a computational model of human speech comprehension. PLoS Biol 21:e3002046.

Takegata R, Morotomi T. 1999. Integrated neural representation of sound and temporal features in human auditory sensory memory: an event-related potential study. Neurosci Lett 274:207–210.

Todd J, Petherbridge A, Speirs B, Provost A, Paton B. 2018. Time as context: The influence of hierarchical patterning on sensory inference. Schizophr Res 191:123–131.

Todorovic A, Auksztulewicz R. 2021. Dissociable neural effects of temporal expectations due to passage of time and contextual probability. Hear Res 399:107871.

Vrba J, Robinson SE. 2001. Signal processing in magnetoencephalography. Methods 25:249–271.

Wilson B, Kikuchi Y, Sun L, Hunter D, Dick F, Smith K, Thiele A, Griffiths TD, Marslen-Wilson WD, Petkov CI. 2015. Auditory sequence processing reveals evolutionarily conserved regions of frontal cortex in macaques and humans. Nat Commun 6:8901.

Wipf D, Nagarajan S. 2009. A unified Bayesian framework for MEG/EEG source imaging. Neuroimage 44:947–966.

Yabe H, Tervaniemi M, Reinikainen K, Näätänen R. 1997. Temporal window of integration revealed by MMN to sound omission. Neuroreport 8:1971–1974.

Ylinen S, Huuskonen M, Mikkola K, Saure E, Sinkkonen T, Paavilainen P. 2016. Predictive coding of phonological rules in auditory cortex: A mismatch negativity study. Brain Lang 162. doi:10.1016/j.bandl.2016.08.007

Zoefel B, Archer-Boyd A, Davis MH. 2018. Phase Entrainment of Brain Oscillations Causally Modulates Neural Responses to Intelligible Speech. Curr Biol 28:401–408.e5.

